# Cell cycle modulates CXCR4 expression in germinal center B cells

**DOI:** 10.1101/175802

**Authors:** Tom S Weber

## Abstract

Adaptation of antibody-mediated immunity occurs in germinal centers (GC). It is where affinity maturation, class switching, memory and plasma cell differentiation synergize to generate specific high affinity antibodies that help both to clear and protect against reinfection of invading pathogens. Within GCs, light and dark zone are two compartments instrumental in regulating this process, by segregating T cell dependent selection and differentiation from generation of GC B cells bearing hypermutated antigen receptors. Spatial segregation of GC B cells into the two zones relies on the chemokine receptor CXCR4, with textbook models attributing high and low expression levels to a dark and light zone phenotype. This bipolarity is however not reflected in the CXCR4 expression profile of GC B cells, which is unimodal and markedly heterogeneous, indicating a continuum of intermediate CXCR4 levels rather than a binary dark or light zone phenotype. Here analysis of published BrdU pulse-chase data reveals that throughout cell cycle, average CXCR4 expression in GC B cells steadily increases by up to 75%, scaling with cell surface area. CXCR4 expression in recently divided GC B cells in G0/G1 phase shows intermediate levels compared to cells in G2 and M phase, consistent with their smaller size. The least number of CXCR4 receptors are displayed by GC B cells in G0/G1 that have not been in cell cycle for several hours. The latter, upon entering S phase however, ramp up relative CXCR4 expression twice as much as recently divided cells. Twelve hours after the BrdU pulse, labelled GC B cells, while initially in S phase, are fully desynchronized in terms of cell cycle and match the CXCR4 expression of unlabeled cells. A model is discussed in which CXCR4 expression in GC B cell increases with cell cycle and cell surface area, with highest levels in G2 and M phase, coinciding with GC B cell receptor signaling in G2 and immediately preceding activation-induced cytidine deaminase (AID) activity in early G1. In the model, GC B cells compete for immobilized or expressed CXCL12 on the basis of their CXCR4 expression levels, gaining a relative advantage as they progress in cell cycle, but loosing the advantage at the moment they divide.

## Introduction

Germinal centers (GC) play a fundamental role in adaptive humoral immunity by providing the niche in which antigen specific activated B cells undergo class switching, affinity maturation, memory and plasma cell differentiation [1–3]. GCs develop in secondary lymphoid organs a few days post immunization or infection. Founded by 20-200 activated B cell clones each [4, 5], they exponentially grow in size, to form a relatively stable broadly sized population [6] and wane several weeks post immunization or after the infection is cleared.

Mature GCs contain GC B cells, T helper cells, tingible body macrophages, a network of follicular dendritic [7] and CXCL12-expressing reticular cells [8]. Each cell type is assigned a specific function in what is collectively termed the germinal center reaction. B cells, as potential effector cells, play a chief part. They generate large amounts of progeny with altered B cell receptors via intense proliferation and activation-induced cytidine deaminase (AID) dependent somatic hypermutation [9, 10]. Some of the progeny undergo AID dependent class-switching [11] and/or division-linked differentiation into memory [12] and long-lived plasma cells [13, 14], while others undergo apoptosis [15]. Most memory cells are derived early [16] while plasma cells are generated late in the response [17]. Key in this complex cell fate decision program are T helper cells that provide survival signals to high affinity GC B cell variants at the expense of lower affinity peers [3, 18–21]. Tingible body macrophages engulf apoptotic GC B cells and debris through phagocytosis and have been proposed to play a role in down-regulating the GC reaction [22]. Follicular dendritic cells stock and supply opsonized antigen coated on their surface via the Fc-receptor [23–25]. Finally, reticular cells produce CXCL12 [8, 26], the ligand for CXCR4, a chemokine receptor essential in polarizing GCs into the light zone (LZ) and dark zone (DZ) [27].

The DZ and LZ are two histologically well defined regions within mature GCs [27–30]. In the DZ, GC B cells divide more frequently [31], and AID, the enzyme required for somatic hypermutation and antibody class switching, is upregulated [32]. In the LZ, follicular dendritic cell network carry and present antigen in form of iccosomes [33], while T helper cells (crucial in providing survival signals to GC B cells), apoptotic cells and tingible body macrophages are more abundant. Taken together these and other observations have led to a model in which GC B cells bearing hypermutated and/or switched antigen receptors are generated primarily in the DZ, and antibody-affinity dependent selection of GC B cells is more likely to happen in the LZ [1, 3]. Implicitly, this model assumes some degree of recycling between the DZ and LZ, the importance of which has long been a matter of debate [34–37].

A major advance in the understanding of cell migration within GCs came with the advent of two-photon live microscopy and live imaging [30, 38, 39]. Monitoring GCs in lymph nodes in anesthetized mice shows highly motile GC B cell, crawling on FDC networks in the LZ, with frequent but mostly short interactions between B and T cells [21, 38, 39]. Due to the limited imaging time windows, precise flux rates between the DZ and LZ have been challenging to infer with this experimental system [40, 41]. To overcome this limitation and quantify cell migration from DZ to LZ over longer time frames an elegant experimental system was developed in which photoactivatable GFP expressing GC B cells were activated in situ in anesthetized mice in either dark or light zone and their position recorded several hours later [42]. This confirmed substantial fluxes between the two zones [43], in line with theoretical predictions developed earlier [36, 37, 44].

The organism level outcome of the GC reaction is affinity maturation [45–47], the increase in average binding affinity of circulating antibodies. Typically described akin to Darwinian evolution [48–51], it involves rounds proliferation and mutation of the genes coding for the B cell receptor variable region (predominantly in the DZ) followed by selection of higher affinity variants and clearance of GC B cells carrying non-functional or low affinity receptors (predominantly in the LZ). While the process by which B cells mutate their BCRs is relatively well understood at the molecular level [52, 53], the details regarding the selection process remain controversial [36, 54–56]. Historically perceived as highly efficient [57], some studies including recent work based on a stochastic multicolor Aid-Cre reporter mice suggests that selection is less stringent than initially thought [5, 58, 59]. Whether low stringency aids in maintaining polyclonality and hence antibody diversity [60], or represents the highest level achievable under biological conditions awaits to be elucidated. Irrespective of the degree of selection pressure however, consensus is that signals from T helper cells are the limiting factor for GC B cell survival [18, 19].

In this work, published BrdU pulse-chase data of GC B cells is reanalyzed [39]. In a first section, proportions of pulse-labelled BrdU+ cells are tracked over time in order to infer turn-over and survival rates of recently divided GC B cells. In a second section, CXCR4 expression is compared between subpopulations that differ in their cell cycle position and DNA content. This reveals the main finding of this study that GC B cell CXCR4 expression steadily increases throughout cell cycle. In two subsequent sections, pulse-chase data at additional time points after the BrdU pulse are analysed. This leads to the identification of two distinct G0/G1 populations that differ in average CXCR4 expression: a first population that has recently divided with intermediate levels, and a second population that has not been in cycle for several hours with low levels. In the last section, analysis of data from cells harvested twelve hours after the pulse shows BrdU labelled and unlabeled cells have converged at that time in terms of CXCR4 expression and cell cycle distribution.

## Materials and Methods

### Experimental procedures

Immunization of mice, BrdU pulse chase and staining procedures are detailed in [39]. In brief, B6 mice were immunized subcutaneously with (4-hydroxy-3-nitrophenyl)acetyl-chicken gamma globulin (NP30- CGG, Biosearch Technologies) emulsified in complete Freund’s adjuvant (Sigma-Aldrich) at 7 different sites. Two weeks later, BrdU (Sigma-Aldrich or BD Pharmingen) in PBS was administered by a single intraperitoneal injection and cells from a total of ten mice were harvested at 30 minutes, 2, 3.5, 5, 8, and 12 hours after the pulse (Fig. S1 A). Draining lymph nodes were pooled for the analysis. BrdU and DNA content was determined using the FITC BrdU flow kit (BD Pharmingen). DAPI was added prior to FACS. For down-stream analysis GC B cells were defined as CD4-CD19+ Fas+ IgDlow cells (Fig. S1 B).

### Analysis of BrdU pulse-chase data

Interpretation of pulse-chase data is based on the relationship between cell cycle progression, cell division and BrdU/DNA content. Fig. S1 C-F shows BrdU and DNA content for three ‘cells’ as they evolve over time after the pulse. For illustrative purposes we assume a clockwise cell cycle progression with G0/G1, S1, S2, S3, S4, S5, G2M as discrete cell cycle states of identical duration (arbitrarily set to 1 hour) and no death (Fig. S1 B). A more realistic model would take into account different phase duration, biologically variability in cell cycle progression and apoptosis [61].

In Fig. S1, initially one of the three cells is in mid-S phase, and has therefore incorporated BrdU (orange cell in Fig. S1 C). A second cell is in G0/G1 phase, and will enter S phase shortly after (red cell). A third cell is in G2M phase (blue cell). Both cells in G0/G1 and G2M are not labelled by BrdU as they are not synthesizing DNA at the time of the pulse. After 3 hours (Fig. S1 D), the BrdU+ cell has reached G2M phase, the cell initially in G0/G1 phase has progressed to mid-S phase, and the cell initially in G2M has divided and its progeny are in early S phase. One hour later (Fig. S1 E), the BrdU+ cell has divided as well, and progeny are in G0/G1 phase. Their BrdU content is half compared to the mother cell and distinguishable from non-labelled cells. The three unlabeled cells have increased their DNA content further, and therefore have ‘moved’ to the right along the DNA axis.

When comparing the schematics in Fig. S1 C-F to real data, there are several additional complexities that need to be considered: a) initially cells are desynchronized, i.e. are distributed all over the cell cycle, b) cell cycle phase duration are variable (e.g. G0/G1 is typically longer than G2M phase), c) cell cycle progression has a significant stochastic component, d) the same cells are not tracked over time, and e) cells undergo apoptosis or differentiate. Despite these differences the simplified model, whose sole purpose is illustration, follows a similar logic, and therefore reflects to a certain degree the underlying dynamics of real bivariate BrdU-DNA content scatter plots after the BrdU pulse.

### Statistical analysis

Error bars in the graphs as well as confidence intervals reported corresponds to mean ± two standard deviations throughout the text. For statistical significance of the difference in means between two samples, the Welch two sample t-test was applied, with p-values < 0.05 being regarded as statistically significant. For linear regression, p-values were computed using the F-test with null hypothesis of the slope being equal zero. Statistical significance between coefficient of variations was tested using the Feltz and Miller test [62] and implemented in the R package ‘cvequality’. All statistical tests were performed using the computing environment R.

## Results

### Pulse-chasing GC B cells confirms high turn-over rates and reveals survival times longer than five hours after birth

GC B cells are highly proliferative, a feature likely to be critical in keeping pace with rapidly evolving pathogens and/or in producing high affinity antibodies most promptly after onset of infection. Consistent with this hypothesis and previous studies [26, 30, 54, 63, 64], in the present data, 24±0.04% GC B cells (compared to 0.063±0.006% in follicular B cells) incorporates the thymidine analog BrdU, which implies that about one in four GC B cells is replicating DNA at any time during the experiment. Although GC B cells are only a zminority (2.4±1%) in draining lymph nodes (LN), they represent 77±0.01% of cells in synthesis (S) phase (Fig. 1 A-B).

**Fig. 1:**
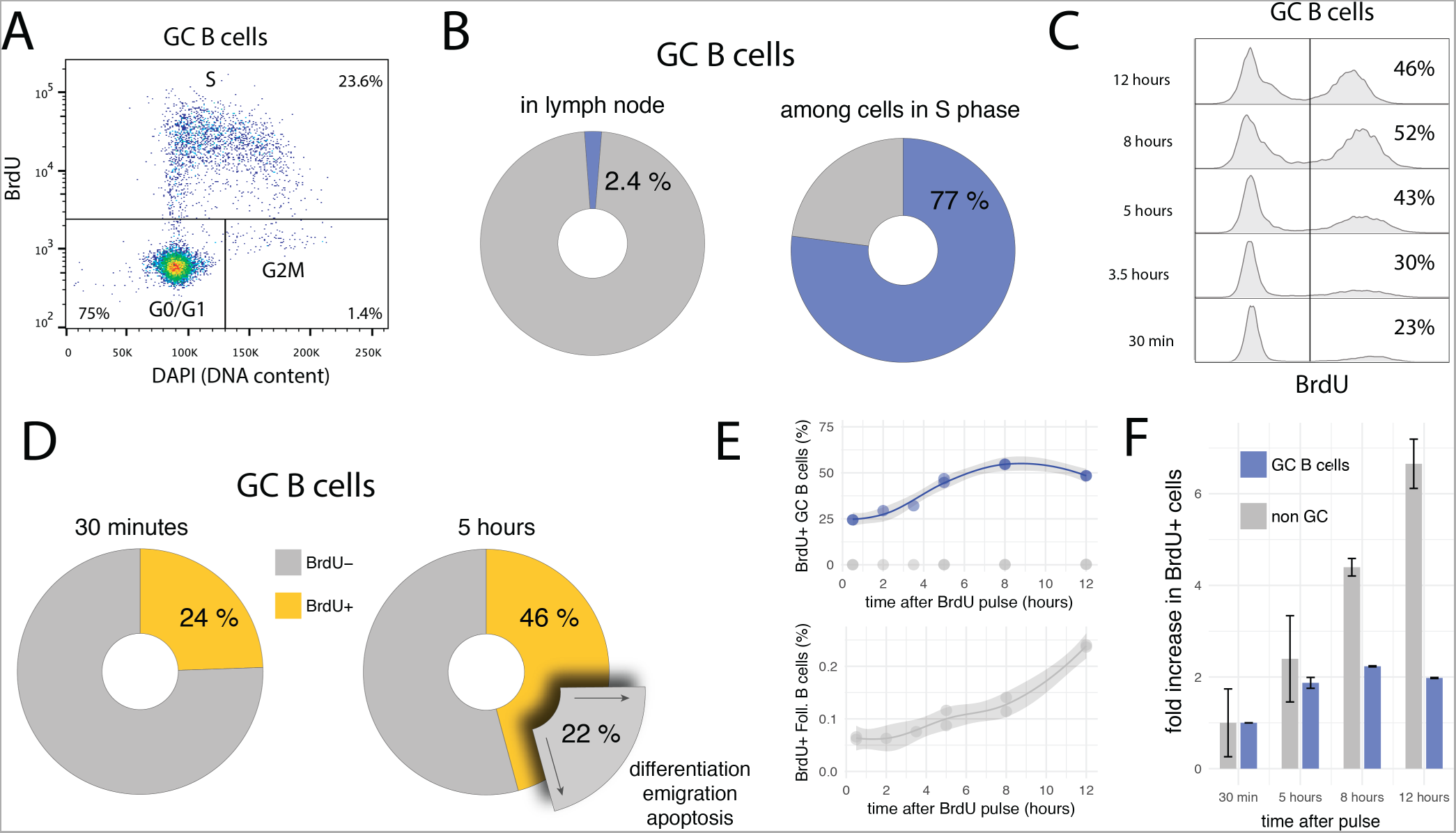
BrdU pulse-chasing of GC B cells. **A)** Bi-variate scatter plot of DAPI (DNA content) versus BrdU 30 minutes after the pulse identifies cells in G0/G1 phase, in S phase, and in G2M phase. **B)** Only a small fraction of cells (2.4±1%) in draining lymph nodes are GC B cells, but due to their fast cycling, they dominate the population of cells in S phase (77±0.01%). **C)** Representative BrdU-pulse chase data, in which GC B cells are harvested at different times (from 30 minutes up to 12 hours) after the BrdU pulse. **D)** Between 30 minutes and 5 hours after the BrdU pulse, the proportion of BrdU+ among GC B cells increases in average by 22%. Assuming quasisteady state (during 4.5 hours), an equivalent proportion of GC B cells must egress from the GC through differentiation or apoptosis during the same period of time. **E-F)** Proportions of BrdU+ GC B cells approximately double within 5-6 hours, and then stabilize at approx. 50%, two-fold of the initial value. In contrast cells that don’t express the GC B cell marker (e.g. follicular B cells) initially contain few BrdU+ cells, but continuously increase this proportion over time.

Some quantitative and qualitative deductions in terms of proliferation and selection/differentiation can be made by analyzing the kinetics of BrdU pulse-labelled cells over time (pulse-chase). Because a single BrdU+ mother cell gives rise to two BrdU+ daughter cells, if the daughter cells survive and maintain a GC B cell phenotype, the proportion of BrdU+ cells has to increase, as soon as labelled cells in late S phase complete G2 and M phase and undergo division. If subsequent survival is longer than the duration of S phase, a doubling in the frequency is expected. Indeed, after a short initial delay, the proportion of BrdU+ GC B cells increases, and almost doubles from 24±0.04% to 46±2% within the first five hours (Fig. 1 C and D). As is readily confirmed on bi-variate FACS plots similar to the one shown in Fig.1 A, this increase is due to labeled cells having divided once and not new cells incorporating BrdU.

The kinetics of labelled cells early after the BrdU pulse are informative in several regards. They indicate that: i) S and G2M phases lasts for about five to six hours, ii) many GC B cells survive and maintain a GC phenotype at least five to six hours after their birth (otherwise the frequency in labelled cells could not increase by a factor close to two), iii) the overall turn-over rate (cell entering cell cycle) in GCs is approximately 25% in five to six hours (in line with 4% of BrdU+EdU-cells per hour observed in recent measurements using the EdU/BrdU double pulse labelling approach, [54] supplementary figure S3). A turn-over rate of 4% per hour is remarkable: In steady state it implies that within six hours, one in four GC B cells either undergoes apoptosis or leaves the GCs via differentiation and emigration (Fig. 1 D). Moreover, because of point ii), many of these cells have not been in cell cycle recently.

A comparison of the relative increase in frequencies of BrdU+ cells in GC B and other LN cell population after the pulse shows a marked difference in kinetics (Fig. 1 E-F). While the level of BrdU+ cells among GC B cells quickly rises and then remains relatively stable at two-fold of the initial value, frequencies of BrdU+ cells in the non-GC B cell populations steadily increase. This difference most likely reflects tight regulation and selection within GCs, as well as constant egress out of the GC. Indeed when BrdU+ cells in non-GC populations are analyzed for DNA content, they mostly appear in G0/G1 phase, therefore possibly representing GC emigrants which have downregulated GC expression markers and have stopped cycling (not shown).

### GC B cell’s CXCR4 expression increases continuously throughout cell cycle

Despite mediating the segregation of GCs into two histologically well defined zones (i.e. light zone and dark zone), CXCR4 expression profile in GC B cells is heterogeneous and does not show two distinguishable peaks (Fig. 2 A), as would be expected from a mixture of high expressing DZ (centroblasts) and low expressing LZ cells (centrocytes). This suggests that CXCR4 expression profiles of centroblasts and centrocytes cells overlap, a feature shared with many other genes expressed in DZ and LZ GC B cells [32]. An exception (published in 2010 after the present data was generated [42]) is CD86, an accessory protein that plays a key role in T cell-B cell costimulation. Together with CXCR4, CD86 is currently the method of choice to gate LZ and DZ GC B cells by flow cytometry (e.g. [18, 54, 64–67]). Nevertheless, some questions regarding CXCR4 expression heterogeneity remain unanswered. For instance, do there exist two or more distinct CXCR4 expression levels in GC B cells that are blurred by stochastic noise or other sources of biological variability? And how is CXCR4 expression related to cell cycle and B cell receptor affinity?

**Fig. 2:**
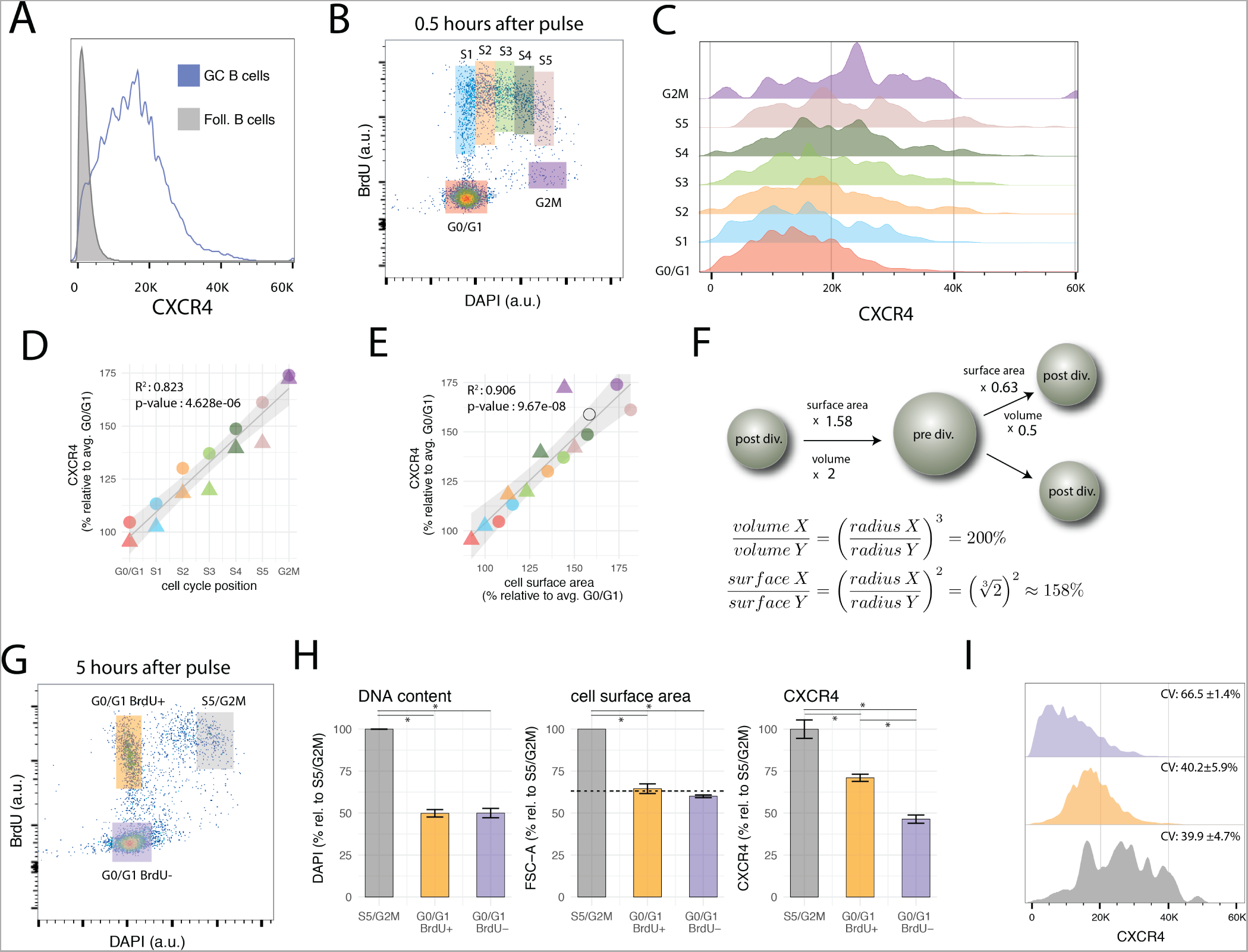
Cell cycle modulates CXCR4 expression in GC B cells. **A)** CXCR4 expression in GC B cells is heterogeneous and its distribution is uni-modal. CXCR4 expression profile of follicular B cells is shown as a negative reference. **B)** Gates used to group GC B cells according to cell cycle position. **C)** CXCR4 expression profile of GC B cells shifts to higher values as cells progress in cell cycle. **D)** Average CXCR4 expression level increases with cell cycle position. Color and shape of the points indicate gate and mouse respectively. **E)** Average CXCR4 expression level increases linearly with cell surface area (as measured by forward scatter). The open circle indicates the relative increase in cell surface area expected from a doubling in volume of a perfect sphere. **F)** Relative increase in surface area of a perfect sphere as it doubles/halves its volume, derived using basic calculus **G)** Representative example of bi-variate scatter plot of DAPI (DNA content) versus BrdU 5 hours after BrdU pulse. Recently divided GC B cells are in G0/G1 BrdU+ gate (orange), while cells that were in S phase during the pulse, and are about to divide are in the S5/G2M gate (grey). GC B cells that have not been in cell cycle in the last 5 hours (and rare cells that were in G2M phase during the pulse) are in the G0/G1 BrdU-gate (violet). **H)** DNA content (DAPI), cell surface area (FSC) and CXCR4 for the gates defined in panel G. The dashed line indicates the reduction in surface area of a perfect sphere that halves its volume. **I)** CXCR4 expression profiles of cells in the S5/G2M, G0/G1 BrdU+, G0/G1 BrdU-gates. As cells divide, they pass from S5/G2M to the G0/G1 BrdU+ gate, resulting in a reduction of average CXCR4 expression by a factor of 0.71±0.01 (panel H), but no change in CV. The population with the highest CV (66.5±1.4%) are the cells in the G0/G1 BrdU-gate, having a low mean but a relatively large spread towards higher levels.

In the present data, BrdU incorporation, DNA content and CXCR4 expression level have been recorded simultaneously with GC phenotypic markers. This permits determination of cell cycle position and CXCR4 expression level in GC B cells after the pulse. Grouping GC B cells into seven gates according to cell cycle position (G0/G1, S1 to S5 and G2M, mapped to cell cycle position {1, 2*, …,* 7}, Fig. 2 B-C) reveals that CXCR4 expression steadily increases from G0/G1 to S and G2M phase (*R*^2^ = 0.82*, p‒value* = 4.6*e*^−6^) with an average value in G2M reaching approximately 75% above G0/G1levels (Fig. 2 D). When plotted against forward scatter, a proxy for cell surface area, CXCR4 also exhibits a linear relationship (Fig. 2 E, *R*^2^ = 0.90*, p ‒ value* = 9.6*e*^-8^), with aslope not statistically different from 1 (*p ‒ value* = 0.23). Together this argues for an increase in total numbers of CXCR4 receptors but maintenance of a relatively consistent surface density throughout cell cycle.

The change in cell surface area is a necessary consequence of the changes in volume of the cell that occur during cell cycle [68]. While the relative increase/decrease depends on the precise shape, the increase in surface of a perfect sphere that doubles its volume is 58% (open circle in Fig. 2 E and Fig 2. F). When the sphere is split into two equally sized smaller spheres, the volume of each is halved, but surface areas are reduced by a factor of 0.63 only (Fig. 2 F). As demonstrated in the next section, CXCR4 expression levels on GC B cells follow a similar trend.

### Low CXCR4 receptor expression of GC B cells in G0/G1 that have not been in cell cycle recently

With CXCR4 expression increasing as cells approach G2M phase, the next question to ask is what happens to the receptors when GC B cells divide. One can address this question (to a certain degree) by comparing, several hours after the BrdU pulse, CXCR4 expression of BrdU+ cells that have just divided, with those that are about to divide (Fig. 2 G). Perhaps as anticipated from our previous results, this analysis shows that recently divided GC B cells in G0/G1 display in average lower numbers of CXCR4 receptors on their surface than their undivided peers in S5/G2M (Fig. 2 H). Several scenarios can be envisioned: for instance CXCR4 receptors are equally aportioned to the daughter cells (dilution), one of the daughter cells receives the majority of the receptors (asymmetric division), or CXCR4 receptor levels are continuously adjusted to cell’s surface area. A reduction by a factor 0.71±0.01 not significantly different from the reduction in cell surface area by 0.64±0.07 (*p‒value* = 0.22), and almost identical coefficients of variation (CV) in CXCR4 expression between cells in S5/G2M and G0/G1 BrdU+ (*p ‒ values* for each mouse are {0.76, 0.96}, Fig. 2 I), argue against dilution (which would result in 50% reduction) or asymmetric aportioning (which would result in a higher CV) of the receptors to the two daughter cells, but suggest an actively regulated process, that maintains cell surface density of CXCR4 receptors approximately constant. Of note is that an identical CV between S5/G2M and G0/G1 BrdU+ cells is expected if daughter cells inherit CXCR4 expression levels proportional to the mother cell (i.e if a mother with relatively high/low CXCR4 expression and surface area generates daughter cells with relatively high/low CXCR4 expression and surface area).

When recently divided cells are compared to BrdU- cells in G0/G1 phase (mostly cells or progeny of cells that are not and have not been in S phase in the last five hours), the CXCR4 expression of the latter is significantly lower, although DNA content and surface area are not (Fig. 2 H, the dashed line indicates the reduction in surface area of a perfect sphere that halves its volume). The BrdU-G0/G1 population is thus enriched for relatively quiescent cells in the LZ, possibly undergoing selection or differentation.

In summary, the above observations demonstrate a continuum of states in terms of CXCR4 expression levels between G0/G1 and M phase, and at least two distinct G0/G1 GC B cell populations with intermediate and low CXCR4 expression levels that differ in their age (or time since last division) and probably location within the GC.

### CXCR4 expression kinetics are different in recently divided and older GC B cells as they reenter cell cycle

We have identified two G1/G0 GC B cell populations that differ in terms of their average CXCR4 profile: recently divided BrdU+ with intermediate and BrdU- cells that have not been in cell cycle recently with low expression levels respectively. What remains unclear is how these two populations evolve over time and how they are related to each other. One possible scenario could be that some time after birth every cell further down-regulates CXCR4 and migrates to the light zone, consistent with a model in which a selection step in the light zone occurs within each division cycle. Such a behavior would be reflected by a decrease in CXCR4 expression in the BrdU+ G0/G1 cell population, prior of entering S phase. Analysis of the present data indicates that this is not the case.

Both recently divided BrdU+ and BrdU-GC B cells in G0/G1 maintain their CXCR4 expression levels. As the cells enter cell cycle, CXCR4 levels increase in both populations (Fig. 3 A-B). Recently divided cells reach a plateau in terms of average CXCR4 copy numbers in mid-S phase (at approximately 50% above G0/G1 levels). In contrast, BrdU- cells, which had been in G0/G1 several hours prior of entering S phase, incessantly ramp up average expression, leading to a two-fold increase in CXCR4 receptors at the end of the cell cycle (Fig. 3 B).

**Fig. 3:**
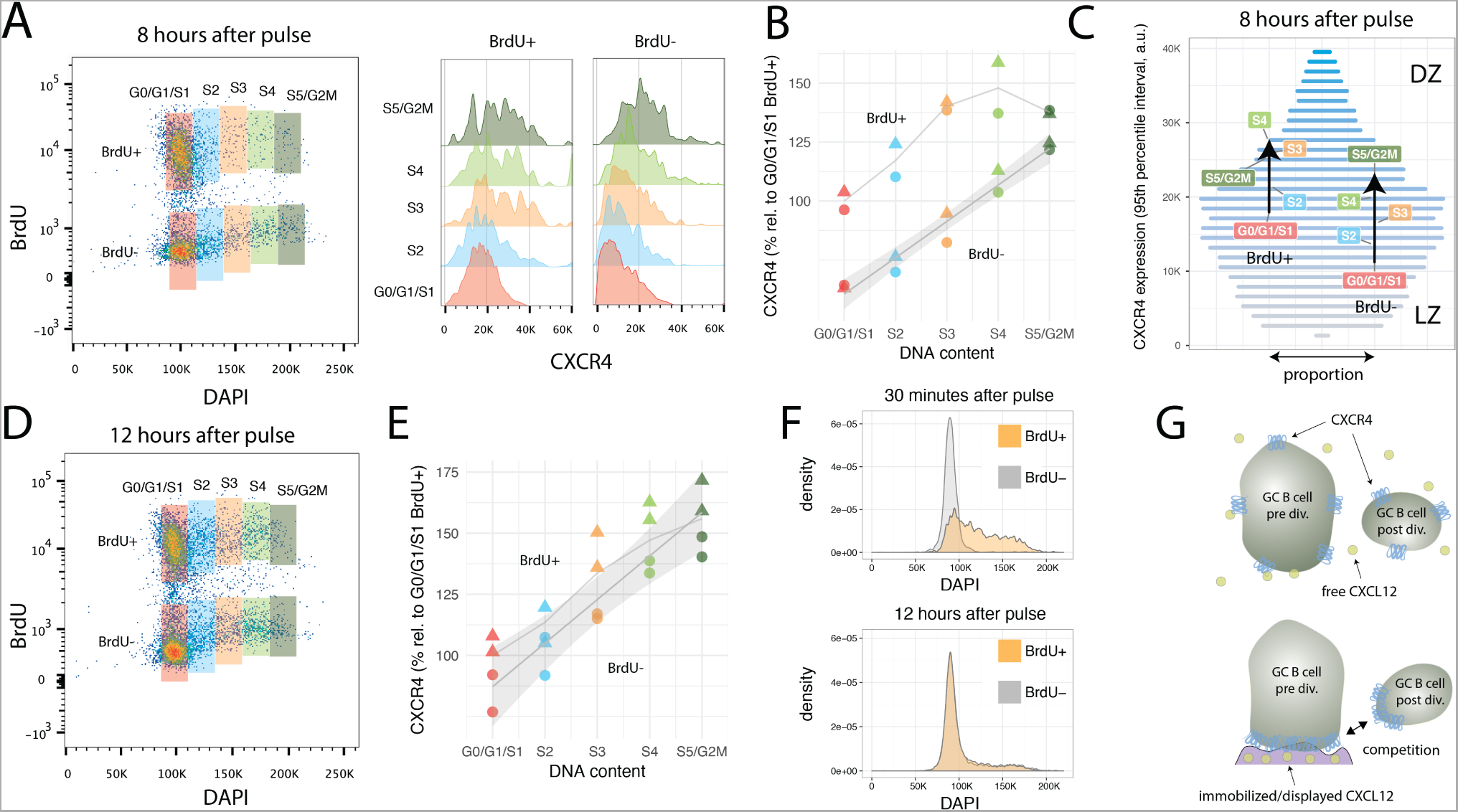
CXCR4 kinetics 8 and 12 hours after BrdU pulse. **A)** Bi-variate scatter plot of DAPI (DNA content) versus BrdU 8 hours after the pulse and corresponding CXCR4 expression profiles. **B)** Average CXCR4 expression increases with cell cycle position in both recently divided BrdU+ and older BrdU- cells. While BrdU+ cells reach a plateau in mid-S at about 50% above their original value, BrdU- cells’ increase in average CXCR4 expression is unabated until G2M phase. Color and shape of the points indicate gate and mouse respectively. **C)** Changes in average CXCR4 expression compared to a representative CXCR4 expression profile. Bar widths correspond to the proportion of GC B cells with a specific CXCR4 expression level (color and y-axis position). DZ and LZ labels indicate that high and low CXCR4 expression is typically attributed to DZ and LZ phenotype respectively. **D)** Bi-variate scatter plot of DAPI (DNA content) versus BrdU 12 hours after the pulse. **E)** CXCR4 expression in the BrdU+ and BrdU-subpopulation has converged indicating complete spatial mixing of individual GCs within approximately 12 hours. Color and shape as in panel B. **F)** Comparison of cell cycle distribution (DNA profile) of BrdU+ and BrdU- cells 30 minutes and 12 hours after pulse. **G)** Proposed theoretical model in which immobilized or displayed CXCL12 leads to competition between cells prior and post division due to their difference in total numbers of CXCR4 receptors. Crucial to this model is the polarization of CXCR4 to the cell’s leading edge in the presence of CXCL12, which has been reported for T cells and various tumor cell lines, but remains to be shown for GC B cells.

Fig. 3 C illustrates how the subpopulations defined in Fig. 3 A are positioned in terms of CXCR4 expression relative to the overall GC B cell population. The widths of the horizontal bars corresponds to the proportion of GC B cells with a given CXCR4 expression level, while their color (and y-axis position) is proportional to the expression level (low: gray, high: blue). For clarity, cells with extreme low and high CXCR4 expression outside the 95% percentiles were excluded from this analysis. BrdU- G0/G1 cells, as they enter and advance in cell cycle, traverse approximately one third of the 95% expression interval largely ‘overtaking’ recently divided cells in G0/G1. The latter however when in mid-S phase reach slightly higher average values, but then stagnate and almost coincide with BrdU- cells in S5/G2M. With the current data, it was not possibly to distinguish whether the reduction in slope represents a general behavior (ie all recently divided GC B cells as they reenter cell cycle are following this trend) or whether some cells downregulate and other cells keep upregulating CXCR4 expression.

### Desynchronization of CXCR4 expression and cell cycle in BrdU+ GC B cells twelve hours after pulse

Twelve hours after the BrdU pulse, CXCR4 levels in BrdU+ are no longer distinguishable from BrdU- cells (Fig. 3 D and E). Similarly, DNA profiles of BrdU+ and BrdU- cells are practically identical (Fig. 3 F). This is remarkable, as most clones probably underwent only one and maximally two divisions since the pulse. Such a rapid desynchronization is indicative for a high variability in GC B cell cycle progression speed, a phenomena perhaps linked to the selection process or diversity in affinity of hypermutated B cell receptors [18].

## Discussion

In this paper, published BrdU pulse-chase GC B cell data from draining lymph nodes two weeks after NP-CCG immunization is reanalyzed. Turn-over rates (cells entering cell cycle) of 4% per hour are inferred for GCs B cells, not inconsistent with GC B cells dividing in average every 12 hours. Despite this fast turn-over, most newly divided cells are found to ‘survive’ for over 5-6 hours after their birth, a time in which hypermutated cells are likely to undergo selection required for affinity maturation.

The analysis further reveals, as its major finding, a so far unreported but potentially far-reaching relationship between GC B cell CXCR4 expression and cell cycle. Average numbers of CXCR4 receptors per cell scale linearly both with DNA content and cell surface area. Compared to BrdU labelled cells in G2M, recently divided BrdU+ GC B cells in G0/G1 displaytimes less CXCR4 receptors on their surface. Expression is further reduced in unlabeled GC B cells in G0/G1, which as witnessed by their BrdU-free DNA, have not been in cell cycle recently. Twelve hours after the pulse, BrdU labelled cells are indistinguishable from unlabeled cells both in terms of cell cycle and CXCR4 expression, suggesting a complete mixing of DZ and LZ in individual GCs within half a day. On a descriptive level, the data demonstrates a greater complexity in GC B cell CXCR4 expression than has previously been appreciated, by linking CXCR4’s heterogeneous expression profile to cell cycle progression and cell division.

What are the implications of the above observations on a theoretical model of the germinal center reaction. While its has long been known that cells in DZ and LZ differ in terms of cell cycle kinetics [28.31, 63], the present analysis suggest that it is cell cycle progression and cell division itself that drives CXCR4 expression and therefore migration towards and against the CXCL12 gradient. The proposed model based on this (and other) data is as follows: As GC B cells progress through cell cycle, surface as well as CXCR4 expression increase. Assuming that CXCR4 in GC B cells polarizes on the cell’s leading edge in the presence of its ligand CXCL12, as has been reported for T cells and several CXCR4 expressing cancer cell lines [69, 70], higher absolute numbers of CXCR4 receptors entail an advantage to compete for space on CXCL12 presenting reticular cell networks (or immobilized CXCL12 on other surfaces) in the DZ (Fig. 3 G). At cell division, total CXCR4 expression drops, as does surface area, in the two daughter cells. This leads to their displacement from the reticular cell network (or other CXCL12 coated surfaces) by other cells at later stages of the cell cycle with higher CXCR4 levels. As a result, the two daughter cells are being ‘pushed back’ towards the LZ consistent with the observed net flux from DZ to LZ [3, 39, 40, 42]. Some GC B cells reenter cell cycle rapidly and start increasing CXCR4 expression levels again (observed in the present data 8 hours after the pulse), while others remain in G0/G1 and decrease CXCR4 further (deduced from the observation that labelled and non labelled cells mix within 12 hours). A proportion of cells with low CXCR4 expression levels that dwell in G0/G1 phase for several hours, reenter S phase, to reach CXCR4 expression levels in G2M similar to cells that reenter S phase immediately after their birth. CXCR4 low expressors in G0/G1 that don’t reenter cell cycle are prone to leave the GC (low levels of CXCR4 could aid these cells to ‘escape’ the CXCR12 gradient) as memory or plasma cells or undergo apoptosis, while intermediate expressors are expanding clones that don’t require T cell help for a further rounds of division.

The data analysed here shows a strong correlation between cell cycle and CXCR4 expression. The proposed model assumes that it is cell cycle that drives CXCR4, but one could argue that it may as well be CXCR4 that regulates cell cycle instead. Evidence that this is most likely not the case comes from a CXCR4 knock out study [26] (similar results have been reported for Foxo1 knock-outs [64–66]), which demonstrates that the lack of expression of this gene does not have a major effect on the magnitude of the GC reaction nor the proportions of cells with light and dark zone phenotype. Intriguingly however, in contrast to proliferation and differentiation, affinity maturation is impaired in these mice. Thus, it seems that GC B cell depend on the CXCR4/CXCL12 axis for effective selection of higher affinity hypermutated variants. How could this relate to cell cycle modulation of CXCR4 levels? Perhaps higher CXCR4 expression levels in G2M ensure that crucial processes like BCR signaling in G2 [71], cytokinesis and AID activity in early G1 phase [72] happen preferentially in proximity to CXCL12, either on the expressing reticular cell network [8] or CXCL12 immobilized on other surfaces [67]. On the other hand, high CXCR4 expression could also enhance BCR signalling in G2 phase, relying on the recently discovered tight link between CXCR4 and BCR ID in mature B cells [73]. Finally increased CXCR4-CXCL12 interaction strength could also potentially facilitate asymmetric cell division [74] by inducing polarization (as reported during T cell development [75]), although in the present data there was no evidence for asymmetric aportioning of CXCR4 to daughter cells. Further studies are needed to answer these questions.

Similar to the CXCR4 knockout, it was shown that for CXCL12^*gagtm*^ mice, in which CXCL12 is unable to bind cellular or extracellular surfaces, magnitude of the germinal center reactions is normal but affinity maturation is less effective [67]. Two observations reported in this study are particularly relevant here. A first one is that GC B cross-section cell surface areas are heterogeneous but significantly larger in DZ then in LZ. A second one is that CXCL12^*gagtm*^ GC B cells in G2M phase are found almost as frequently in LZ as in DZ while in wild-type controls the majority is found in the DZ only. Both observations are in line with the model proposed above in which a CXCL12 gradient serves as a guide for cycling cells to reach CXCL12 high regions when approaching G2M phase.

The weakness (and perhaps strength) of this work is the small number of samples it is based on (i.e. 10 mice in total) and the fact that the data was created using a single experimental technique (i.e. BrdU pulse-chase). Clearly the hypotheses generated by this study remain to be challenged in future experiments. Repeats with different immunization protocols, timings, and mouse strains will help to test the robustness of the observed relationships and kinetics. And additional surface markers, for instance Ki-67 to separate G0 and G1 cells, Blimp-1 to identify plasma blasts [16], and/or the recently discovered marker Ephrin-B1 which marks mature GC B cells [76], will aid to further resolve the fate, cell cycle and CXCR4 expression levels of relevant subpopulations. Technically more involved approaches, for instance continuously monitoring CXCR4 expression in cycling GC B cells from CXCR4 cross FUCCI reporter mice, via in vitro long-term imaging and tracking would also be highly informative [77, 78], as would be GC B single-cell RNA sequencing experiments [79].

Beyond its function in affinity maturation, CXCR4 is implicated in regulating numerous other processes, for example embryonic development [80, 81], hematopoietic stem cell self-renewal in the bone marrow [82], and neutrophil release during stress [83]. Its role in disease further highlights its relevance in cellular homing and proliferation. CXCR4 is over expressed in more than 23 human cancers [84] including leukemia [85], is associated with metastasation [86], and has been identified as a marker for poor prognosis in human patients [87]. For HIV it represents a major cofactor for entry into T-cells during the immunological deficient phase of infection [88]. If cell cycle modulates CXCR4 expression in GC B cell, as the data analysed here indicates, it will be important to investigate whether this mechanism is specific to GCs or whether it also plays a role in other tissues and cell types.

## ACKNOWLEDGMENTS

Christopher Allen for generously sharing his data, Phil Hodgkin for critically reading and commenting on the manuscript, Michal Or-Guil and Jorge Carneiro for many insightful discussions on germinal centers.

---

**Fig. S1:**
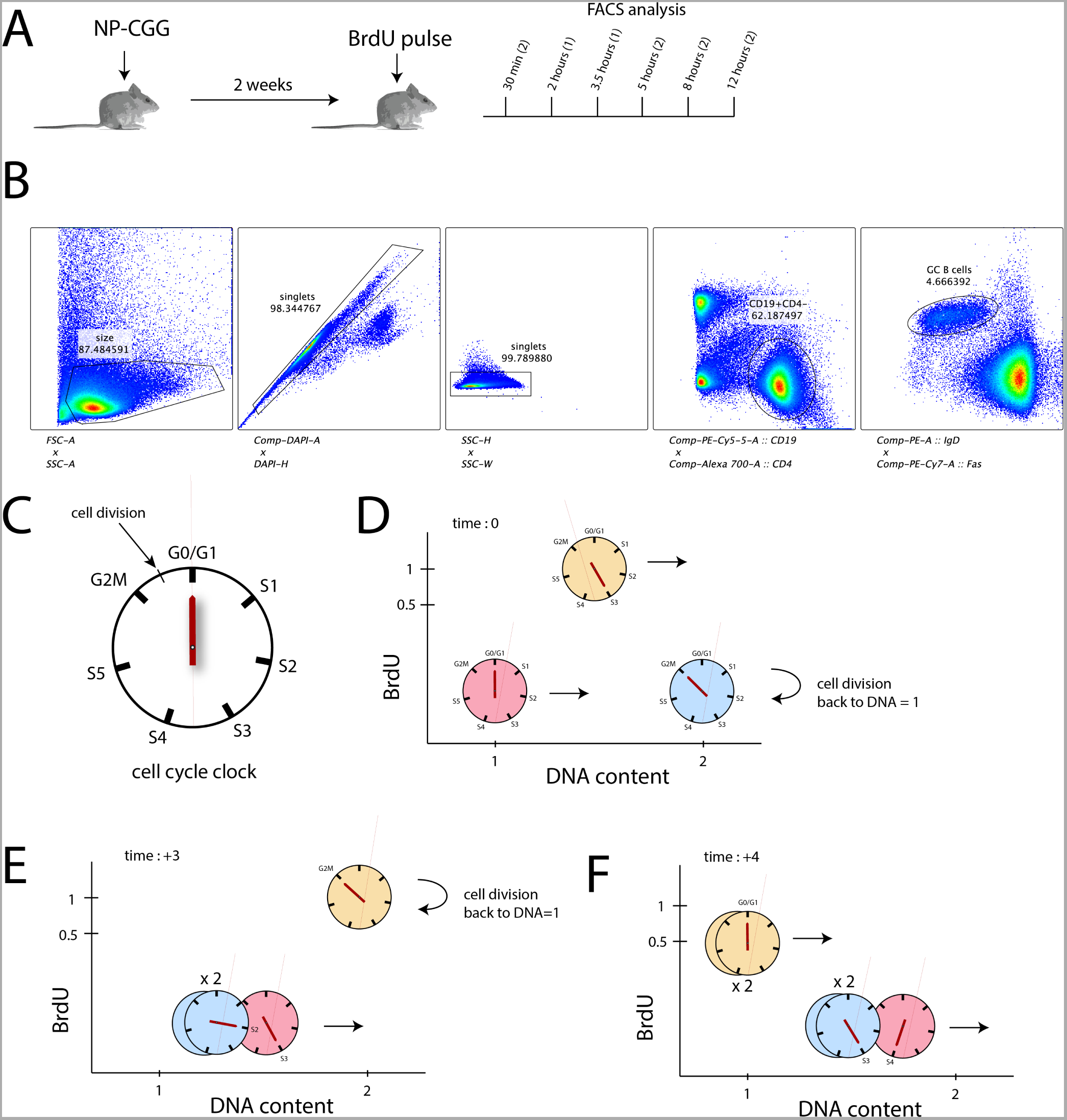
Experimental setup and interpretation of BrdU pulse-chase data. **A)** Overview of immunization and BrdU labeling protocol. Numbers in brackets give the number of mice sampled at each time point. **B)** Gating strategy used to isolate single GC B cells. **C)** Simplified cell cycle clock model with G0/G1, S1, S2, S3, S4, S5, and G2M as discrete phases. For the purpose of illustration all phases are assumed to have the same duration (1 hour) and cell division occurs between G2M and G1. **D)** Three virtual cells in different phases of the cell cycle immediately after a BrdU pulse. Cell cycle position determines position of the DNA versus BrdU plot. As cells continue replication they ‘move’ to the right as DNA content increases, eventually divide, and ‘jump back’ to DNA content 1 at division. **E)** Three hours after the pulse, the BrdU labelled cell in panel D has reached G2M phase, the cell initially in G0/G1 is half-way through S phase, and the cell initially in G2M has divided and its progeny are in early S phase. **F)** Four hours after the pulse, the BrdU+ cell has divided as well, and the three unlabeled cells have moved further to the right.

## References

1. DeFranco, A. L. (2016) The germinal center antibody response in health and disease. F1000Research 5, 99+.

2. Klein, U & Dalla-Favera, R. (2008) Germinal centres: role in B-cell physiology and malignancy. Nature reviews. Immunology 8, 22–33.

3. Allen, C. D, Okada, T, & Cyster, J. G. (2007) Germinal-center organization and cellular dynamics. Immunity 27, 190–202.

4. Faro, J & Or-Guil, M. (2013) How oligoclonal are germinal centers? A new method for estimating clonal diversity from immunohistological sections. BMC bioinformatics 14 Suppl 6, S8+.

5. Tas, J. M. J, Mesin, L, Pasqual, G, Targ, S, Jacobsen, J. T, Mano, Y. M, Chen, C. S, Weill, J.-C, Reynaud, C.-A, Browne, E. P, Meyer-Hermann, M, & Victora, G. D. (2016) Visualizing antibody affinity maturation in germinal centers. Science pp. aad3439+.

6. Wittenbrink, N, Weber, T. S, Klein, A, Weiser, A. A, Zuschratter, W, Sibila, M, Schuchhardt, J, & Or-Guil, M. (2010) Broad Volume Distributions Indicate Nonsynchronized Growth and Suggest Sudden Collapses of Germinal Center B Cell Populations. The Journal of Immunology 184, 1339–1347.

7. Allen, C. D & Cyster, J. G. (2008) Follicular dendritic cell networks of primary follicles and germinal centers: phenotype and function. Seminars in immunology 20, 14–25.

8. Rodda, L. B, Bannard, O, Ludewig, B, Nagasawa, T, & Cyster, J. G. (2015) Phenotypic and Morphological Properties of Germinal Center Dark Zone Cxcl12-Expressing Reticular Cells. The Journal of Immunology pp. 1501191+.

9. Muramatsu, M, Sankaranand, V. S, Anant, S, Sugai, M, Kinoshita, K, Davidson, N. O, & Honjo, T. (1999) Specific Expression of Activation-induced Cytidine Deaminase (AID), a Novel Member of the RNA-editing Deaminase Family in Germinal Center B Cells. Journal of Biological Chemistry 274, 18470–18476.

10. Muramatsu, M, Kinoshita, K, Fagarasan, S, Yamada, S, Shinkai, Y, & Honjo, T. (2000) Class switch recombination and hypermutation require activation-induced cytidine deaminase (AID), a potential RNA editing enzyme. Cell 102, 553–563.

11. Stavnezer, J, Guikema, J. E. J, & Schrader, C. E. (2008) Mechanism and Regulation of Class Switch Recombination. Annual Review of Immunology 26, 261–292.

12. Tarlinton, D. (2006) B-cell memory: are subsets necessary? Nature Reviews Immunology 6, 785–790.

13. Nutt, S. L & Tarlinton, D. M. (2011) Germinal center B and follicular helper T cells: siblings, cousins or just good friends? Nature immunology 12, 472–477.

14. Hasbold, J, Corcoran, L. M, Tarlinton, D. M, Tangye, S. G, & Hodgkin, P. D. (2004) Evidence from the generation of immunoglobulin G-secreting cells that stochastic mechanisms regulate lymphocyte differentiation. Nature immunology 5, 55–63.

15. Li, Y, Takahashi, Y, Fujii, S.-i, Zhou, Y, Hong, R, Suzuki, A, Tsubata, T, Hase, K, & Wang, J.-Y. (2016) EAF2 mediates germinal centre B-cell apoptosis to suppress excessive immune responses and prevent autoimmunity. Nature Communications 7, 10836+.

16. Blink, E. J, Light, A, Kallies, A, Nutt, S. L, Hodgkin, P. D, & Tarlinton, D. M. (2005) Early appearance of germinal center-derived memory B cells and plasma cells in blood after primary immunization. The Journal of experimental medicine 201, 545–554.

17. l, F. J, Zuccarino-Catania, G. V, Chikina, M, & Shlomchik, M. J. (2016) A Temporal Switch in the Germinal Center Determines Differential Output of Memory B and Plasma Cells. Immunity 44, 116–130.

18. Gitlin, A. D, Mayer, C. T, Oliveira, T. Y, Shulman, Z, Jones, M. J. K, Koren, A, & Nussenzweig, M. C. (2015) T cell help controls the speed of the cell cycle in germinal center B cells. Science 349, 643–646.

19. Meyer-Hermann, M, Maini, P. K, & Iber, D. (2006) An analysis of B cell selection mechanisms in germinal cen-ters.

20. De Silva, N. S & Klein, U. (2015) Dynamics of B cells in germinal centres. Nature Reviews Immunology 15, 137– 148.

21. Shulman, Z, Gitlin, A. D, Weinstein, J. S, Lainez, B. n, Esplugues, E, Flavell, R. A, Craft, J. E, & Nussenzweig, M. C. (2014) Dynamic signaling by T follicular helper cells during germinal center B cell selection. Science 345, 1058–1062.

22. Smith, J. P, Burton, G. F, Tew, J. G, & Szakal, A. K. (1998) Tingible body macrophages in regulation of germinal center reactions. Developmental immunology 6, 285– 294.

23. Kosco-Vilbois, M. H. (2003) Are follicular dendritic cells really good for nothing? Nature Reviews Immunology 3, 764–769.

24. Tew, J. G, Wu, J, Fakher, M, Szakal, A. K, & Qin, D. (2001) Follicular dendritic cells: beyond the necessity of T-cell help. Trends in Immunology pp. 361–367.

25. Vora, K. A, Ravetch, J. V, & Manser, T. (1997) Amplified follicular immune complex deposition in mice lacking the Fc receptor gamma-chain does not alter maturation of the B cell response. J Immunol 159, 2116–2124.

26. Bannard, O, Horton, R. M, Allen, C. D. C, An, J, Nagasawa, T, & Cyster, J. G. (2017) Germinal Center Centroblasts Transition to a Centrocyte Phenotype According to a Timed Program and Depend on the Dark Zone for Effective Selection. Immunity 39, 912–924.

27. Allen, C. D, Ansel, K. M, Low, C, Lesley, R, Tamamura, H, Fujii, N, & Cyster, J. G. (2004) Germinal center dark and light zone organization is mediated by CXCR4 and CXCR5. Nature immunology 5, 943–952.

28. MacLennan, I. C. (1994) Germinal centers. Annual review of immunology 12, 117–139.

29. Camacho, S. A, Kosco-Vilbois, M. H, & Berek, C. (1998) The dynamic structure of the germinal center. Immunology today 19, 511–514.

30. Hauser, A. E, Junt, T, Mempel, T. R, Sneddon, M. W, Kleinstein, S. H, Henrickson, S. E, von Andrian, U. H, Shlomchik, M. J, & Haberman, A. M. (2007) Definition of germinal-center B cell migration in vivo reveals predominant intrazonal circulation patterns. Immunity 26, 655–667.

31. Hanna, M. G. (1964) An autoradiographic study of the germinal center in spleen white pulp during early intervals of the immune response. Laboratory investigation; a journal of technical methods and pathology 13, 95–104.

32. Victora, G. D, Dominguez-Sola, D, Holmes, A. B, Deroubaix, S, Dalla-Favera, R, & Nussenzweig, M. C. (2012) Identification of human germinal center light and dark zone cells and their relationship to human B cell lymphomas. Blood 120, blood–2012–03–415380–2248.

33. Suzuki, K, Grigorova, I, Phan, T. G, Kelly, L. M, & Cyster, J. G. (2009) Visualizing B cell capture of cognate antigen from follicular dendritic cells. Journal of Experimental Medicine 206, 1485–1493.

34. Hanna, M. G, Congdon, C. C, & Wust, C. J. (1966) Effect of antigen dose on lymphatic tissue germinal center changes. Proceedings of the Society for Experimental Biology and Medicine. Society for Experimental Biology and Medicine (New York, N.Y.) 121, 286–290.

35. Moreira, J. S & Faro, J. (2006) Re-evaluating the recycling hypothesis in the germinal centre. Immunology and Cell Biology 84, 404–410.

36. Oprea, M & Perelson, A. S. (1997) Somatic mutation leads to efficient affinity maturation when centrocytes recycle back to centroblasts. Journal of immunology (Baltimore, Md. : 1950) 158, 5155–5162.

37. Meyer-Hermann, M. (2001) Recycling Probability and Dynamical Properties of Germinal Center Reactions. Journal of Theoretical Biology 210, 265–285.

38. Schwickert, T. A, Lindquist, R. L, Shakhar, G, Livshits, G, Skokos, D, Kosco-Vilbois, M. H, Dustin, M. L, & Nussenzweig, M. C. (2007) In vivo imaging of germinal centres reveals a dynamic open structure. Nature 446, 83–87.

39. Allen, C. D. C, Okada, T, Tang, H. L, & Cyster, J. G. (2007) Imaging of Germinal Center Selection Events During Affinity Maturation. Science 315, 528–531.

40. Beltman, J. B, Allen, C. D. C, Cyster, J. G, & de Boer, R. J. (2011) B cells within germinal centers migrate preferentially from dark to light zone. Proceedings of the National Academy of Sciences 108, 8755–8760.

41. Meyer-Hermann, M. E & Maini, P. K. (2005) Interpreting two-photon imaging data of lymphocyte motility. Physical review. E, Statistical, nonlinear, and soft matter physics 71.

42. Victora, G. D, Schwickert, T. A, Fooksman, D. R, Kamphorst, A. O, Meyer-Hermann, M, Dustin, M. L, & Nussenzweig, M. C. (2010) Germinal center dynamics revealed by multiphoton microscopy with a photoactivatable fluorescent reporter. Cell 143, 592–605.

43. Meyer-Hermann, M, Mohr, E, Pelletier, N, Zhang, Y, Victora, G. D, & Toellner, K.-M. M. (2012) A theory of germinal center B cell selection, division, and exit. Cell reports 2, 162–174.

44. Kepler, T. B & Perelson, A. S. (1993) Cyclic re-entry of germinal center B cells and the efficiency of affinity maturation. Immunology today 14, 412–415.

45. Berek, C, Berger, A, & Apel, M. (1991) Maturation of the immune response in germinal centers. Cell 67, 1121–1129.

46. Tarlinton, D. (2000) Dissecting affinity maturation: a model explaining selection of antibody-forming cells and memory B cells in the germinal centre. Immunology Today 21, 436–441.

47. Takahashi, Y, Dutta, P. R, Cerasoli, D. M, & Kelsoe, G. (1998) In situ studies of the primary immune response to (4-hydroxy-3-nitrophenyl)acetyl. V. Affinity maturation develops in two stages of clonal selection. The Journal of experimental medicine 187, 885–895.

48. Dunn-Walters, D. K, Belelovsky, A, Edelman, H, Banerjee, M, & Mehr, R. (2002) The Dynamics of Germinal Centre Selection as Measured by Graph-Theoretical Analysis of Mutational Lineage Trees. Developmental Immunology pp. 233–243.

49. Steiman-Shimony, A, Edelman, H, Hutzler, A, Barak, M, Zuckerman, N. S, Shahaf, G, Dunn-Walters, D, Stott, Mehr, R. (2006) Lineage tree analysis of immunoglobulin variable-region gene mutations in autoimmune diseases: Chronic activation, normal selection. Cellular Immunology 244, 130–136.

50. Kleinstein, S. H, Louzoun, Y, & Shlomchik, M. J. (2003) Estimating Hypermutation Rates from Clonal Tree Data. J Immunol 171, 4639–4649.

51. Tarlinton, D. M. (2008) Evolution in miniature: selection, survival and distribution of antigen reactive cells in the germinal centre. Immunol Cell Biol 86, 133–138.

52. Peled, J. U, Kuang, F. L, Iglesias Ussel, M. D, Roa, S, Kalis, S. L, Goodman, M. F, & Scharff, M. D. (2008) The Biochemistry of Somatic Hypermutation. Annual Review of Immunology 26, 481–511.

53. Di Noia, J. M & Neuberger, M. S. (2007) Molecular Mechanisms of Antibody Somatic Hypermutation. Annual Review of Biochemistry 76, 1–22.

54. Gitlin, A. D, Shulman, Z, & Nussenzweig, M. C. (2014) Clonal selection in the germinal centre by regulated proliferation and hypermutation. Nature 509, 637–640.

55. Liu, Y. J, Joshua, D. E, Williams, G. T, Smith, C. A, Gordon, J, & Maclennan, I. C. M. (1989) Mechanism of antigen-driven selection in germinal centres. Nature 342, 929–931.

56. Meyer-Hermann, M. E & Maini, P. K. (2005) Cutting edge: back to "one-way" germinal centers. J Immunol 174, 2489–2493.

57. Kepler, T. B & Perelson, A. S. (1993) Somatic hypermutation in B cells: an optimal control treatment. J Theor Biol 164, 37–64.

58. Kleinstein, S. H & Singh, J. P. (2003) Why are there so few key mutant clones? The influence of stochastic selection and blocking on affinity maturation in the germinal center. Int. Immunol. 15, 871–884.

59. Radmacher, M. D, Kelsoe, G, & Kepler, T. B. (1998) Predicted and inferred waiting times for key mutations in the germinal centre reaction: Evidence for stochasticity in selection. Immunol Cell Biol 76, 373–381.

60. Greiff, V, Bhat, P, Cook, S. C, Menzel, U, Kang, W, & Reddy, S. T. (2015) A bioinformatic framework for immune repertoire diversity profiling enables detection of immunological status. Genome Medicine 7.

61. Weber, T. S, Jaehnert, I, Schichor, C, Or-Guil, M, & Carneiro, J. (2014) Quantifying the Length and Variance of the Eukaryotic Cell Cycle Phases by a Stochastic Model and Dual Nucleoside Pulse Labelling. PLOS Computational Biology 10, e1003616+.

62. Feltz, C. J & Miller, G. E. (1996) An asymptotic test for the equality of coefficients of variation from k populations. Statistics in medicine 15, 646–658.

63. Liu, Y. J, Zhang, J, Lane, P. J, Chan, E. Y, & MacLennan, ndary responses to T cell-dependent and T cell-independent antigens. European journal of immunology 21, 2951–2962.

64. Sander, S, Chu, V. T, Yasuda, T, Franklin, A, Graf, R, Calado, D. P, Li, S, Imami, K, Selbach, M, Di Virgilio, M, Bullinger, L, & Rajewsky, K. (2017) PI3 Kinase and FOXO1 Transcription Factor Activity Differentially Control B Cells in the Germinal Center Light and Dark Zones. Immunity 43, 1075–1086.

65. Dominguez-Sola, D, Kung, J, Holmes, A. B, Wells, V. A, Mo, T, Basso, K, & Dalla-Favera, R. (2017) The FOXO1 Transcription Factor Instructs the Germinal Center Dark Zone Program. Immunity 43, 1064–1074.

66. Inoue, T, Shinnakasu, R, Ise, W, Kawai, C, Egawa, T, & Kurosaki, T. (2017) The transcription factor Foxo1 controls germinal center B cell proliferation in response to T cell help. Journal of Experimental Medicine pp. jem.20161263+.

67. Barinov, A, Luo, L, Gasse, P, Meas-Yedid, V, Donnadieu, E, Arenzana-Seisdedos, F, & Vieira, P. (2017) Essential role of immobilized chemokine CXCL12 in the regulation of the humoral immune response. Proceedings of the National Academy of Sciences 114, 2319–2324.

68. Marguerat, S & Bähler, J. (2012) Coordinating genome expression with cell size. Trends in Genetics 28, 560–565.

69. Gómez-Moutón, C, Abad, J. L, Mira, E, Lacalle, R. A, Gallardo, E, Jiménez-Baranda, S, Illa, I, Bernad, A, Mañes, S, & Carlos Martínez, A. (2001) Segregation of leading-edge and uropod components into specific lipid rafts during T cell polarization. Proceedings of the National Academy of Sciences 98, 9642–9647.

70. van Buul, J. D, Voermans, C, van Gelderen, J, Anthony, E. C, van der Schoot, C. E, & Hordijk, P. L. (2003) Leukocyte-Endothelium Interaction Promotes SDF-1-dependent Polarization of CXCR4. Journal of Biological Chemistry 278, 30302–30310.

71. Khalil, A. M, Cambier, J. C, & Shlomchik, M. J. (2012) B Cell Receptor Signal Transduction in the GC Is Short-Circuited by High Phosphatase Activity. Science 336, 1178–1181.

72. Wang, Q, Kieffer-Kwon, K.-R, Oliveira, T. Y, Mayer, C. T, Yao, K, Pai, J, Cao, Z, Dose, M, Casellas, R, Jankovic, M, Nussenzweig, M. C, & Robbiani, D. F. (2016) The cell cycle restricts activation-induced cytidine deaminase activity to early G1. Journal of Experimental Medicine pp. jem.20161649+.

73. Becker, M, Hobeika, E, Jumaa, H, Reth, M, & Maity, P. C. (2017) CXCR4 signaling and function require the expression of the IgD-class B-cell antigen receptor. Proceedings of the National Academy of Sciences 114, 5231– 5236.

74. Barnett, B. E, Ciocca, M. L, Goenka, R, Barnett, L. G, Wu, J, Laufer, T. M, Burkhardt, J. K, Cancro, M. P, & Reiner, S. L. (2012) Asymmetric B Cell Division in the Germinal Center Reaction. Science 335, 342–344.

75. Pham, K, Shimoni, R, Charnley, M, Ludford-Menting, M. J, Hawkins, E. D, Ramsbottom, K, Oliaro, J, Izon, D, Ting, S. B, Reynolds, J, Lythe, G, Molina-Paris, C, Melichar, H, Robey, E, Humbert, P. O, Gu, M, & Russell, S. M. (2015) Asymmetric cell division during T cell development controls downstream fate. J Cell Biol 210, 933–950.

76. Laidlaw, B. J, Schmidt, T. H, Green, J. A, Allen, C.D. C, Okada, T, & Cyster, J. G. (2017) The Eph-related tyrosine kinase ligand Ephrin-B1 marks germinal center and memory precursor B cells. Journal of Experimental Medicine pp. jem.20161461+.

77. Pauklin, S & Vallier, L. (2013) The Cell-Cycle State of Stem Cells Determines Cell Fate Propensity. Cell 155, 135–147.

78. Kinjyo, I, Qin, J, Tan, S.-Y, Wellard, C. J, Mrass, P, Ritchie, W, Doi, A, Cavanagh, L. L, Tomura, M, Sakaue-Sawano, A, Kanagawa, O, Miyawaki, A, Hodgkin, P. D, & Weninger, W. (2015) Real-time tracking of cell cycle progression during CD8+ effector and memory T-cell differentiation. Nature Communications 6, 6301+.

79. Wu, Y. L, Stubbington, M. J. T, Daly, M, Teichmann, S. A, & Rada, C. (2017) Intrinsic transcriptional heterogeneity in B cells controls early class switching to IgE. Journal of Experimental Medicine 214, 183–196.

80. Pujol, F, Kitabgi, P, & Boudin, H. (2005) The chemokine SDF-1 differentially regulates axonal elongation and branching in hippocampal neurons. J Cell Sci 118, 1071–1080.

81. Raz, E & Mahabaleshwar, H. (2009) Chemokine signaling in embryonic cell migration: a fisheye view. Development 136, 1223–1229.

82. Sugiyama, T, Kohara, H, Noda, M, & Nagasawa, T.(2006) Maintenance of the Hematopoietic Stem Cell Pool by CXCL12-CXCR4 Chemokine Signaling in Bone Marrow Stromal Cell Niches. Immunity 25, 977–988.

83. Eash, K. J, Means, J. M, White, D. W, & Link, D. C. (2009) CXCR4 is a key regulator of neutrophil release from the bone marrow under basal and stress granulopoiesis conditions. Blood 113, 4711–4719.

84. Chatterjee, S, Behnam Azad, B, & Nimmagadda, S. (2014) The Intricate Role of CXCR4 in Cancer. (Elsevier) Vol. 124, pp. 31–82.

85. Burger, J. A & Bürkle, A. (2007) The CXCR4 chemokine receptor in acute and chronic leukaemia: a marrow homing receptor and potential therapeutic target. British journal of haematology 137, 288–296.

86. Muller, A, Homey, B, Soto, H, Ge, N, Catron, D, Buchanan, M. E, McClanahan, T, Murphy, E, Yuan, W, Wagner, S. N, Barrera, J. L, Mohar, A, Verastegui, E, & Zlotnik, A. (2001) Involvement of chemokine receptors in breast cancer metastasis. Nature 410, 50–56.

87. Scala, S, Ottaiano, A, Ascierto, P. A, Cavalli, M, Simeone, E, Giuliano, P, Napolitano, M, Franco, R, Botti, G,CR4 Predicts Poor Prognosis in Patients with Malignant Melanoma. Clinical Cancer Research 11, 1835–1841.

88. Vicenzi, E, Liò, P, & Poli, G. (2013) The Puzzling Role of CXCR4 in Human Immunodeficiency Virus Infection. Theranostics3, 18–25.

